# Photoprotection comes at a cost: non-photochemical quenching shapes growth–productivity trade-offs in *Haematococcus lacustris*

**DOI:** 10.64898/2026.08.19.745743

**Authors:** Stefano Cazzaniga, Francesco Bellamoli, Edoardo Ceschi, Laura Girolomoni, Nico Olivieri, Mattia Magagnotti, Matteo Paloschi, Marzia Rossato, Massimo Delledonne, Matteo Ballottari

## Abstract

Non-photochemical quenching (NPQ) dissipates excess absorbed light energy and protects photosynthetic organisms from photodamage, but its role in regulating the balance between growth, stress tolerance and astaxanthin accumulation in *Haematococcus lacustris* remains unclear. Here, we investigated how enhanced NPQ affects photosynthetic performance, stress-induced differentiation, and productivity in this astaxanthin-producing microalga.

We isolated and characterized an NPQ-enhanced mutant line, A116, using cultivation assays under different stress conditions, analysis of photosynthetic parameters, pigment profiling, and whole-genome resequencing. A116 displayed stronger and faster NPQ induction, driven by increased LHCSR accumulation, resulting in decreased photosynthetic electron transport and lower photochemical efficiency under moderate-to-high light. Enhanced NPQ delayed the transition to astaxanthin-rich cysts under high light, allowing greater biomass accumulation under CO_2_-limiting conditions. However, under high CO_2_ availability, where carbon fixation relieved excitation pressure supporting efficient photosynthesis, the enhanced NPQ phenotype reduced growth and astaxanthin productivity compared with the wild type.

These results show that NPQ modulates a context-dependent trade-off between photoprotection and productivity in *Haematococcus lacustris*. Increased energy dissipation can improve high-light tolerance under carbon limitation, but becomes detrimental when absorbed light can be efficiently used for carbon assimilation. Thus, optimal algal productivity requires tuning photoprotective capacity to environmental conditions rather than maximizing NPQ.

**HIGHLIGHT:** In astaxanthin-producing microalgae, enhanced photoprotection improves high-light tolerance and delays stress-induced differentiation, but limits biomass and astaxanthin productivity when carbon availability enables efficient photosynthesis.

## INTRODUCTION

*Haematococcus lacustris* (previously named *pluvialis*) is a freshwater unicellular green microalga, belonging to the Chlorophyceae class that undergoes, under unfavorable conditions, into a transition to peculiar cel stage called cyst (or haematocyst) characterized by a massive accumulation of the ketocarotenoid astaxanthin (3, 3′-dihydroxy-β, β’-carotene-4, 4′-dione) (Boussiba *et al*., 1999). Astaxanthin is a carotenoid with a bright red color used as a coloring agent mainly in aquaculture and poultry (Lorenz and Cysewski, 2000). It is also a strong antioxidant, preventing the production of reactive oxygen species (ROS) and other oxidative molecules (Fan *et al*., 1998; Kobayashi, 2003; Kobayashi and Okada, 2000; Lemoine and Schoefs, 2010; Mascia *et al*., 2017; Scibilia *et al*., 2015; Shah *et al*., 2016; Song *et al*., 2014; Zhang *et al*., 2014). Thanks to this antioxidant activity, astaxanthin has promising effects in the prevention and treatment of several diseases such as cancers, inflammations, cardiovascular and gastrointestinal diseases, and in enhancing the immune system (Capelli *et al*., 2019; Guerin *et al*., 2003; Koyande *et al*., 2019; Yuan *et al*., 2011; Zhang *et al*., 2014). Among the organisms capable of producing astaxanthin, *H. lacustris* can accumulate the largest amount, up to 5-6% of its dry weight (Boussiba, 2000; Boussiba *et al*., 1999; Boussiba and Vonshak, 1991).

Under favorable environmental conditions, *H. lacustris* lives mainly as free-swimming, biflagellated vegetative cells that can asexually reproduce and increase biomass (Boussiba, 2000). In this stage, *H. lacustris* cells are photosynthetically active: the pigment-binding complexes Photosystem I (PSI) and Photosystem II (PSII), embedded in the thylakoid membranes inside the chloroplasts, absorb light energy and convert it into chemical energy used to produce NADPH and ATP. PSII and PSI have similar elements distributed in two different moieties: the antenna and the core complex (Shen *et al*., 2019; Su *et al*., 2019). Antenna proteins absorb light and transfer excitation energy to the reaction center, where charge separation occurs. From the PSII core, electrons pass through plastoquinone, cytochrome b6f complex, and plastocyanin to reach PSI, where the second reaction of charge separation takes place. From PSI, electrons move to ferredoxin and the final acceptor, NADPH. Electron transport through the chain is coupled to the generation of a protonic gradient across the thylakoid membrane, which is utilized by ATP synthase to produce ATP. Together with NADPH, ATP is used in the light-independent phase of photosynthesis to fix CO_2_ into organic molecules. The main pigment bound to PSII and PSI and responsible for light absorption is chlorophyll *a*. Additional pigments that broaden the wavelengths of light absorbed are chlorophyll b and carotenoids. In the vegetative stage of *H. lacustris*, the carotenoids present are β-carotene and the xanthophylls lutein, violaxanthin, zeaxanthin, and neoxanthin, while astaxanthin is not accumulated (Mascia *et al*., 2017).

Under stressful conditions, vegetative cells become round, expand in cell size, and lose both flagella (Kobayashi *et al*., 1991). In this stage, *H. lacustris* forms immobile cysts, protected by a thick cell wall, that do not duplicate and do not accumulate additional biomass (Kobayashi *et al*., 2001). These haematocysts are characterized by exceptional resilience to various adverse conditions, enabling them to withstand extreme environments. Astaxanthin biosynthesis begins during the transition from the vegetative stage to the intermediate non-motile stage, called palmella, and it reaches its maximum after the transition to the mature stage of haematocysts (Scibilia *et al*., 2015; Wang *et al*., 2014). Astaxanthin is mainly accumulated in droplets in the perinuclear cytoplasm (Chen *et al*., 2015; Gwak *et al*., 2014; Ota *et al*., 2018; Wayama *et al*., 2013), resulting in the bright red color of the cells. During the red phase, astaxanthin accounts for more than 80% of the total carotenoids, mainly in its esterified form (Chen *et al*., 2014; Holtin *et al*., 2009). The transition to haematocysts does not affect only the carotenoid composition, but it is a complex process with major changes in cell ultrastructure, metabolism, and chloroplast electron transport chain: transition to astaxanthin-rich cysts leads to a decrease in chlorophyll content, destabilization of the photosynthetic apparatus and inhibition of the photosynthetic activity (Boussiba, 2000; Boussiba *et al*., 1999; Lemoine and Schoefs, 2010; Mascia *et al*., 2017; Scibilia *et al*., 2015; Solovchenko *et al*., 2013). When growth conditions revert to favorable, the hematocyst releases astaxanthin-containing zoospores that later revert to vegetative cells. The transition from “green to red cells” is triggered by different types of stresses like high salinity, high or low temperature, nutrient deprivation, and, in particular, high light (Zhekisheva *et al*., 2002a). The susceptibility to adverse culture conditions leads to a substantial reduction in productivity, posing a major obstacle that negatively influences the industrial performance of *H. lacustris* cultivation. This microalga is sensitive to subtle changes in nutrient and environmental conditions, including pH, light intensity, temperature, and dissolved oxygen, that could limit cell division and biomass accumulation (Hong *et al*., 2015; Kobayashi, 2003; Lemoine and Schoefs, 2010). Various studies are actively conducted to improve astaxanthin productivity in *H. lacustris* (Acheampong *et al*., 2025; Azizi *et al*., 2024; Fábregas *et al*., 2001; Kayani *et al*., 2024; Lopez *et al*., 2006; Morgado *et al*., 2024; Park *et al*., 2014; Peng *et al*., 2025; Zhou *et al*., 2025). One potential target is the trade-off between astaxanthin accumulation and photoprotective mechanisms.

One of the main stresses triggering the transition to the haematocycst is the excessive increase of irradiance. Theoretically, the growth rate of vegetative algae cells increases with increasing light intensity until the photosynthetic apparatus becomes saturated. Excess light energy saturates or unbalances the electron transport chain, resulting in the production of ROS that can react with proteins, lipids, and nucleic acids, generating photodamage (Andersson *et al*., 1992; Niyogi, 1999). Photosynthetic organisms have developed different mechanisms to protect themselves from such damage. The main one, in the short period, is non-photochemical quenching (NPQ) that dissipates the excess light energy absorbed as heat (Niyogi, 1999). This process involves PSII antenna subunits and their bound carotenoids (Ruban *et al*., 2011).

According to the time scale over which it occurs and relaxes in the dark, NPQ can be defined into three major components. The qE is the fastest and most relevant component (1-2 minutes to activate and relax) and depends on the generation of the electrochemical proton gradient across the thylakoid membrane (Pascal *et al*., 2005) being sensed by some specific membrane proteins as PSBS and LHCSR, the former being the main actor for qE in land plants (Fan *et al*., 2015; Li *et al*., 2000b; Li *et al*., 2004) while LHCSR and LHCSR-like proteins (as LHCX in diatoms) are the main players for qE in microalgae (Bailleul *et al*., 2010b; Blommaert *et al*., 2017; Ghazaryan *et al*., 2016; Lepetit *et al*., 2012; Niyogi and Truong, 2013; Peers *et al*., 2009; Steen *et al*., 2022; Zhu and Green, 2010). The second, intermediate component (relaxing within 5-15 minutes in the dark) is named qT or qZ and reflects the state transition of the antenna of the PSII and the xanthophyll cycle (de-epoxidation of violaxanthin into zeaxanthin) (Bellafiore *et al*., 2005; Nilkens *et al*., 2010). The slowest component, qI (hours to relax in the dark), is related to photoinhibition and PSII core D1 subunit turnover (Matsubara and Chow, 2004). NPQ response changes during red phase transition in *H. lacustris,* showing a biphasic pattern (Chekanov *et al*., 2019). In the early phase, there is an increase in quenching, followed by a strong reduction, culminating in the complete suppression of the mechanism in mature haematocysts (Chekanov *et al*., 2016a). While the photosynthetic apparatus begins to disassemble, the cells are more prone to imbalanced high-light fluxes, since chlorophylls are still present and light energy is absorbed, but the electron transport chain is already being dismantled (Scibilia *et al*., 2015). The increase in NPQ during the early transition phase likely mitigates ROS formation in this transitory phase. However, it is important to note that ROS are associated with cellular signaling components leading to stress-dependent astaxanthin biosynthesis in *H. lacustris*. High-light stress and NPQ induction could be closely linked in regulating the transition of *H. lacustris* from green cells to astaxanthin-rich cysts. The relationship between the changes associated with astaxanthin accumulation and photoprotective mechanisms remains unclear and warrants further investigation. The impact of NPQ on productivity is context-dependent in different photosynthetic species: in *Chlamydomonas reinhardtii*, reduced qE capacity and impaired state transitions increase sensitivity to high light (Allorent *et al*., 2016; Allorent *et al*., 2013), whereas in crops, accelerating NPQ relaxation enhances photosynthetic efficiency and biomass production under fluctuating light (Kromdijk *et al*., 2016; Long *et al*., 2025).

In this study, we investigated the role of NPQ in regulating photoprotection, photosynthetic performance, and stress-induced astaxanthin accumulation in *Haematococcus lacustris*. To this aim, we isolated and characterized a mutant strain (A116) displaying enhanced NPQ capacity and examined its physiological responses under high-light conditions and contrasting inorganic carbon availabilities. By combining analyses of photosynthetic activity, photoprotective responses, growth, and astaxanthin production, we assessed how increased NPQ influences the balance between light stress tolerance and productivity. By combining analyses of photosynthetic activity, photoprotective responses, growth and astaxanthin production, we assessed how increased NPQ influences the balance between light stress tolerance and productivity in *H. lacustris*.

## MATERIALS AND METHODS

### Haematococcus lacustris cultivation

The *Haematococcus lacustris* strain K-0084 was obtained from the Scandinavian Culture Collection of Algae and Protozoa. Stock cultures were grown photoautotrophically in BG-11 medium at 22 °C under continuous illumination (25 µmol photons m^-2^ s^-1^) in flasks on a rotary shaker (150 rpm) to ensure proper mixing and gas exchange (Scibilia *et al*., 2015).

Stress conditions to induce ketocarotenoid formation were applied as described below. Exponentially growing cultures (approximately 5 × 10^5^ cells mL^-1^) were subjected to high light, high salinity, nitrogen deprivation, or control conditions (BG-11 medium at 25 µmol photons m^-2^ s^-1^) and cultivated for 7 days in 50 mL flasks at 22°C. High-light conditions were applied at 300 µmol photons m^-2^ s^-1^. High salinity was achieved by supplementing BG-11 medium with 1% NaCl. For nitrogen starvation, nitrate was omitted from the medium; prior to treatment, cells were harvested by centrifugation and washed three times with nitrate-free BG-11. All experiments were performed on a rotary shaker (150 rpm) to prevent sedimentation, with three independent biological replicates per condition. Samples (1 mL) were collected daily for pigment analysis.

Semicontinuous cultivation was performed as follows. *H. lacustris* cultures grown in flasks at 25 µmol photons m^-2^ s^-1^ in BG-11 were diluted to an initial density of 5×10^5^ cell mL^-1^ and exposed to 300 µmol photons m^-2^ s^-1^. After four days, when cultures reached saturation, each flask was diluted 1:3 and grown for an additional three days under the same conditions. This cycle was repeated twice. For additional nitrogen starvation, at the end of the cycle, cells were harvested by centrifugation (1000 g, 5 min), washed with nitrogen-depleted BG-11 (without NaNO_3_), and resuspended in 50 mL of the same medium. Samples were collected before each dilution step and at the end of the experiment. Semicontinuous cultivation was performed on WT and A116 strains with three independent biological replicates per strain.

Cultivation under high CO_2_ availability was performed using a Multi-Cultivator MC1000 system (Photon System Instrument, Czech Republic) equipped with eight airlift photobioreactors (80 mL volume). Cultures were continuously supplied with air enriched with 3% CO_2_ and grown under continuous illumination at 300, 600, 1200 or 3000 µmol photons m^−2^ s^−1^ at 22°C on BG-11 medium.

### Haematococcus lacustris transformation and mutant screening

Transformation of *Haematococcus lacustris* was performed by *Agrobacterium tumefaciens*-mediated insertional mutagenesis using strain EHA105 carrying the binary vector pCAMBIA1300. The vector harbors the hygromycin phosphotransferase gene (*hptII*) under the control of the CaMV 35S promoter, enabling selection of transformants through random T-DNA integration into the nuclear genome. The transformation protocol was adapted from a previous report (Kathiresan *et al*., 2015), as described below.

For transformation, approximately 10⁶ algal cells were pre-cultivated on rich solid medium for 5-7 days. *A. tumefaciens* was grown overnight in LB medium and subsequently expanded in fresh medium to reach an OD₆₀₀ of 1.2. Bacterial cells were then harvested by centrifugation, washed, and resuspended in induction medium consisting of algal rich medium supplemented with 100 μM acetosyringone (pH 5.6) to a final OD₆₀₀ of 1. Algal cells were collected from plates, mixed with 200 μL of *A. tumefaciens* suspension, and spread onto induction agar medium. Co-cultivation was carried out for 2 days to allow T-DNA transfer. Following infection, cells were recovered in liquid rich medium supplemented with carbenicillin (100 mg L^-1^) and incubated for 30 min to eliminate bacterial cells. Algal cells were then collected by centrifugation, washed with BG-11 supplemented with acetate, and plated onto solid selection medium containing hygromycin (25 mg L^-1^) and carbenicillin (100 mg L^-1^). Plates were incubated at 25 °C until the appearance of resistant colonies.

Putative transformants were isolated and maintained on selective medium. Integration of the hygromycin resistance cassette was verified by PCR using *hptII*-specific primers (forward: 5′-GCTGCATCATCGAAATTGCC-3′; reverse: 5′-TTATCGGCACTTTGCATCGG-3′).

Primary transformants were screened for alterations in non-photochemical quenching (NPQ) using a chlorophyll fluorescence imaging system (FluorCam 700MF, Photon Systems Instruments, Brno, Czech Republic). Colonies grown on solid selective medium were initially analyzed directly to identify lines displaying significant deviations in NPQ compared to the wild type. A threshold of ±40% variation relative to WT values was used to select candidate lines with altered quenching capacity. Selected colonies were transferred to liquid medium and grown in the absence of hygromycin prior to a second round of NPQ measurements performed under standardized conditions to confirm the phenotype. To assess the stability of the transformation, selected lines were re-plated on solid medium containing hygromycin. However, non-resistant lines displaying stable NPQ phenotypes were also retained for physiological analyses.

### Pigments analysis

Pigments were extracted from intact cells using DMSO (Zhekisheva *et al*., 2002b) as described in (Scibilia *et al*., 2015). The chlorophyll-to-carotenoid ratio and chlorophyll *a*/*b* ratio were estimated from the absorption spectra of pigment extracts as described in (Perozeni *et al*., 2020). Absorption spectra were recorded using a Jasco V-550 UV-visible spectrophotometer and analyzed by spectral fitting (Chazaux *et al*., 2022). Carotenoid composition was further analyzed by high-performance liquid chromatography (HPLC) as previously described (Perozeni *et al*., 2020).

### Gel electrophoresis and immunobloFng

Non-denaturing Deriphat-PAGE was performed following solubilization of isolated thylakoidal membranes. Thylakoid were purified as previously described (Cazzaniga *et al*., 2014). Samples corresponding to 30 μg of chlorophylls were solubilized with 0.6% (w/v) n-dodecyl-α-D-maltoside (α-DM) and separated using Midi BioRad gel systems (Girolomoni *et al*., 2020). SDS-PAGE was performed using Mini BioRad gel systems loaded with increasing amounts of solubilized thylakoids corresponding to 0.25, 0.5, 1, or 2 μg of chlorophyll. Proteins were transferred to membranes and analyzed by Western blotting as previously described. The following primary antibodies (Agrisera, Sweden) were used: αPsaA (AS06 172), αCP47 (AS04 038), αLHCII (AS01 003), α-LHCSR3 (AS14 2766), α-PSBS (AS06 167), α-CP26 (AS09 407), α-CP29 (AS04 045). Secondary detection was performed using an anti-rabbit IgG antibody (A3687, Merck) conjugated to alkaline phosphatase, with chromogenic detection.

### Oxygen evolution, photosynthetic parameters, and NPQ measurements

Light-dependent oxygen evolution was measured using a Clark-type O_2_ electrode (Oxygraph Plus, Hansatech) at 25°C. Samples with a cell density of (5×10^6^ cells ml^-1^) were transferred to a 1 x 1 cm cuvette mixed by magnetic stirring for oxygen measurements. Dark respiration was measured in the absence of light, while light-dependent oxygen evolution was recorded upon illumination with a halogen lamp (Schott) at different actinic light intensities (50-2500 µmol photons m^−2^ s^−1^) in the presence of 5 mM sodium bicarbonate. Net oxygen evolution rates were obtained by subtracting dark respiration from light-dependent oxygen production and fitted using a hyperbolic function: y = P_max_ * x/(K_i_ + x), where P_max_ is the maximum net oxygen evolution rate and K_i_ the light intensity at which the rate reaches P_max_/2 (Supplementary Table S1). Photosynthetic parameters, including ΦPSII, 1 - qL, ETR, and NPQ (Baker, 2008; Van Kooten and Snel, 1990), were measured using a DUAL-PAM-100 fluorimeter (Heinz-Walz, Germany). Measurements were performed on whole cells at a density of 3 x 10^6^ cells ml^-1^ in a 1×1 cm cuvette under continuous magnetic stirring at room temperature. Samples were exposed to a saturating light pulse (4000 µmol photons m^−2^ s^−1^) and to actinic light intensities ranging from 50 to 1200 µmol photons m^−2^ s^−1^.

### Electrochromic shift

Electrochromic shift (ECS) measurements were performed as previously described (Kuhlgert *et al*., 2016). Briefly, cells (1×10^7^ cells ml^-1^) were dark-adapted for 20 minutes, and ECS was measured in a 2 mm cuvette using a MultispeQ v2.0 instrument (PhotosynQ) at different actinic light intensities (100-1200 μmol photons m^−2^ s^−1^). ECS measurements were performed both in the presence and absence of 50 μM DCMU to inhibit linear electron flow. DCMU was applied to whole cells with a 10 min incubation time prior to measurements.

### DNA extraction

Genomic DNA was extracted as previously described in (Marcolungo *et al*., 2024). Briefly, nuclei were isolated from approximately 4 × 10^8^ *H. lacustris* cells using MEB buffer (Lutz *et al*., 2011), and nuclear DNA was purified using a Genomic-tip-100/G kit (Qiagen, Hilden, Germany). DNA concentration was determined using the Qubit dsDNA BR Assay Kit and a Qubit fluorimeter (Thermo Fisher Scientific). DNA purity was evaluated by spectrophotometric analysis using a NanoDrop instrument (Thermo Fisher Scientific), and DNA integrity was assessed using a TapeStation 4150 system with a Genomic DNA ScreenTape assay (Agilent Technologies).

### Illumina sequencing

Whole-genome sequencing libraries were prepared using the KAPA HyperPrep Kit (Kapa Biosystems) following a PCR-free protocol. Nuclear DNA was sheared using an M220 ultra-sonicator (Covaris), adjusting the treatment time to obtain ∼350 bp fragments. Library size distribution was assessed by capillary electrophoresis using a Bioanalyzer High Sensitivity DNA chip (Agilent Technologies). Libraries were quantified by qPCR using a standard curve and sequenced in 150 bp paired-end mode on an Illumina NovaSeq 6000 platform, generating 45,231,415 read pairs.

### Variant calling and annotation

Sequencing adapters and low-quality bases were trimmed using fastp v0.21. Filtered reads were mapped to the *H. lacustris* reference genome (Marcolungo *et al*., 2024) using BWA-MEM2 v2.2.1 (Vasimuddin *et al*., 2019). Duplicates were identified and removed using GATK v4.1.7.0 (MarkDuplicates), and overlapping regions of paired-end reads were clipped using fgbio v1.3.0 ClipBam tool (http://fulcrumgenomics.github.io/fgbio/). Variant calling was performed on the A116 mutant and WT datasets (Marcolungo *et al*., 2024) using FreeBayes v1.3.6, with ploidy set to 2. Only variants with base quality ≥ 20, mapping quality ≥ 20, and read depth ≥ 10 were retained. Additional filtering was applied using bcftools v1.10.2 (QUAL > 20 and MQM > 49), and variants shared between samples were removed using SnpSift v5.0d. Variant annotation was performed using SnpEff v5.0d (Cingolani *et al*., 2012). The ANN field of the annotated VCF file was parsed to extract predicted functional consequences and impact categories. Variant summaries were generated from three datasets: the complete set of variants, variants overlapping annotated genes, and variants located within coding sequences (CDS). These datasets were obtained by applying genomic feature-based filtering to the annotated VCF. Each dataset was processed independently using the same analysis workflow to ensure consistency across comparisons. Only variants present in the analyzed sample (i.e. genotypes different from 0/0) were considered. Variants were stratified by genotype (e.g. heterozygous 0/1, homozygous 1/1, and multi-allelic genotypes), and all counts were reported separately for each genotype class. Genotype stratification allows distinguishing between heterozygous and homozygous variants, providing additional resolution on the distribution of predicted functional effects. Variant consequences were summarized at the variant level, such that each consequence term was counted at most once per variant, irrespective of the number of annotated transcripts. Combined consequence annotations (e.g. missense_variant&splice_region_variant) were decomposed into their individual components and counted separately. Variants were classified according to their predicted impact (HIGH, MODERATE, LOW, MODIFIER). In addition to transcript-aware impact counts, each variant was also assigned a single maximum impact category based on the hierarchy HIGH > MODERATE > LOW > MODIFIER. For clarity, a subset of major coding consequence categories was reported in the summary (Supplementary Table S2), while a complete list of all detected consequence terms is provided in the complete list (Supplementary Table S3).

### Functional annotation and enrichment analysis of variant-associated genes

The *H. lacustris* genome was functionally re-annotated using eggNOG-mapper v2.1.13 (Cantalapiedra *et al*., 2021) based on orthology assignments obtained with DIAMOND. Functional categories, including Gene Ontology (GO), KEGG pathways, and COG classifications, were retrieved. Functional enrichment analyses were performed in R using the clusterProfiler package (Yu *et al*., 2012). Gene sets were constructed from eggNOG-mapper annotations, and enrichment was tested using a hypergeometric model implemented in the enricher function. The background universe comprised all genes with available eggNOG annotations, and only annotated genes were included in the analysis. The input gene list consisted of non-redundant genes carrying variants identified from the filtered VCF file, restricted to coding sequences (CDS) and to variants with predicted impacts classified as HIGH, MODERATE, or LOW, while variants annotated as MODIFIER were excluded. P-values were adjusted for multiple testing using the Benjamini-Hochberg method, and terms with adjusted p-values < 0.05 were considered significantly enriched. Genes carrying variants but lacking eggNOG annotation were excluded from enrichment analyses but retained for downstream interpretation based on the original genome annotation (Marcolungo *et al*., 2024). A comprehensive table including all variant-carrying genes was generated, reporting for each gene whether it was included in the enrichment analysis (based on the availability of eggNOG annotations) and integrating both eggNOG-mapper functional annotations and the original genome annotations (Supplementary Table S4).

### Statistical analysis

Statistical analyses were performed using two-tailed Welch’s t-tests to compare WT and A116 samples. Data are presented as mean ± SD (standard deviation). Differences were considered statistically significant at p < 0.05.

## RESULTS

### Selection of a Haematococcus lacustris mutant with increased NPQ

Exposure to high light stress was reported to induce an increased NPQ induction in *H. lacustris* (Chekanov *et al*., 2016a; Scibilia *et al*., 2015). To investigate the potential effect of NPQ on the transition from the green to the red phase in *H. lacustris*, an insertional mutagenesis approach was applied to generate mutants with increased NPQ phenotype. Insertional mutagenesis is a well-established method used across various species to generate novel strains that can be selected for desired phenotypes. The experimental design applied to *H. lacustris* involved random insertion of an antibiotic (hygromycin) resistance cassette via *Agrobacterium tumefaciens* infection (Kathiresan *et al*., 2009),followed by screening for strains exhibiting increased NPQ relative to the WT. A vector carrying a hygromycin resistance gene as a selection marker was used. A transformation efficiency of 0.068% was obtained, with 2745 surviving colonies confirmed by replating on selective medium (Fig. 1A). The surviving lines were then screened for NPQ (Fig. 1B): 87 exhibited increased or decreased NPQ relative to the WT, using a 40% variation from the WT as a threshold. These lines were then transferred to liquid medium without hygromycin and rescreened for NPQ using the same 40% variation threshold. While 35 lines were confirmed to have lower NPQ than the WT, only one line, designated A116, showed increased quenching (Fig. 1C). However, when the selected 36 lines were transferred back to solid medium in the presence of the selective marker, only 12 lines were resistant, whereas the remaining 24 lines failed to survive, indicating the absence of an integrated hygromycin resistance cassette in their genome. The line exhibiting increased NPQ (A116) was among those that did not survive on the selective medium; nevertheless, due to its interesting phenotype, the presence of the antibiotic cassette was further investigated by PCR. As shown in Supplementary Fig. S1, the hygromycin resistance cassette was indeed lost in A116, whereas it was retained in some of the other lines analyzed.

**Figure 1.**
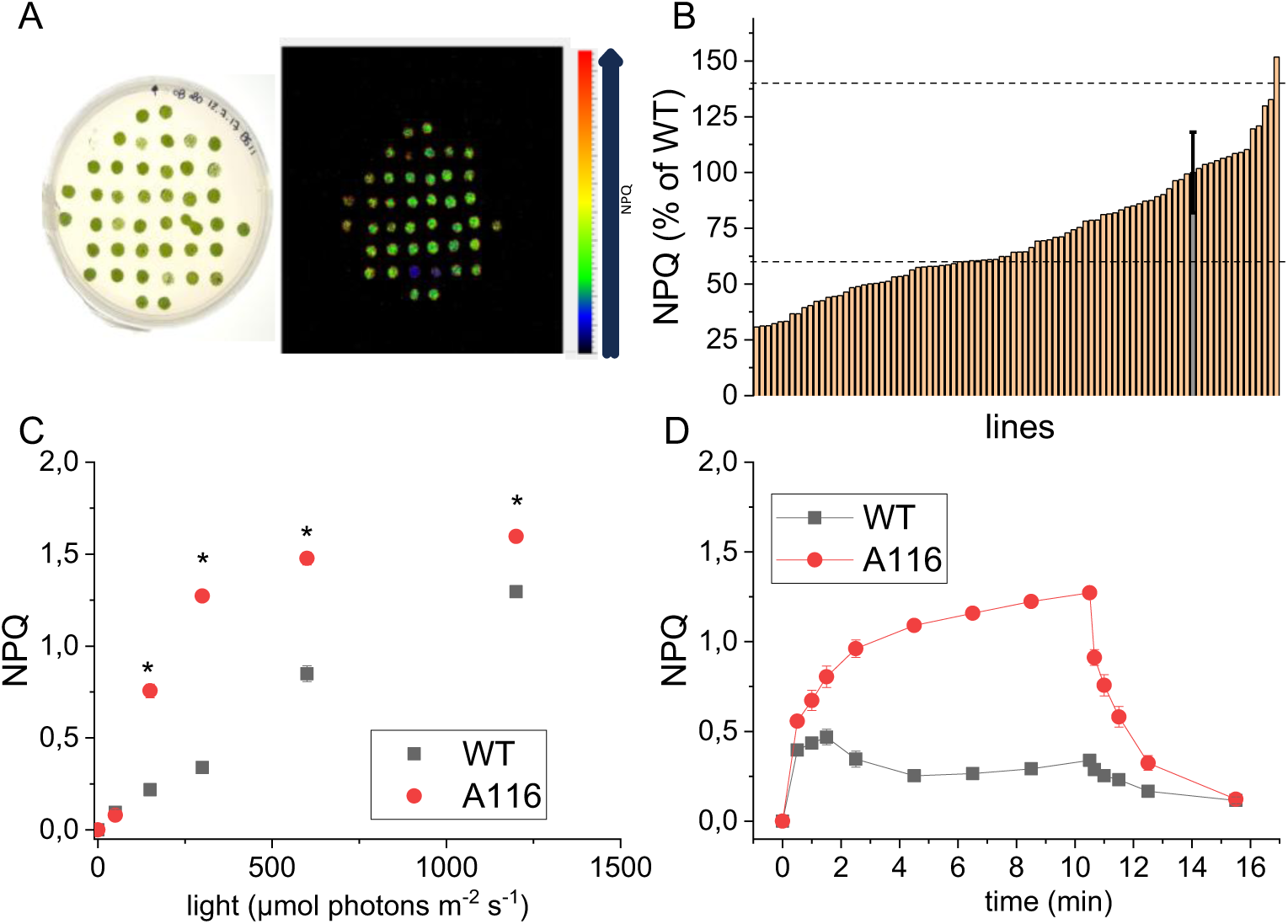
Generation and screening of *H. lacustris* insertional mutants. (A) Representative agar plate for NPQ screening using FluorCam 700MF (left) and instrument output (right). (B) NPQ screening of *H. lacustris* lines. WT is represented as a gray bar, while dotted lines represent 40% threshold respect to WT NPQ. (C) NPQ measured on wild type (WT, black) and A116 (red) cells using different actinic lights. Values correspond to NPQ at the end of the 10 minutes illumination. Data are presented as mean ± SD (n = 4). * indicate significant differences between WT and A116 (p < 0.05). (D) NPQ measured on WT and A116 cells using an actinic light of 300 μmol photons m^-2^ s^-1^. ALT TEXT: Panel A shows representative agar plates and fluorescence imaging output used to screen *Haematococcus lacustris* mutant colonies for non-photochemical quenching. Panel B shows the primary NPQ screening of mutant lines, with wild type values and threshold lines indicating 40% variation from the wild type. Panel C shows NPQ values measured under increasing actinic light intensities in wild type and A116 cells, with A116 displaying higher NPQ from 150 to 1200 μmol photons m^-2^ s^-1^. Panel D shows NPQ induction kinetics at 300 μmol photons m^-2^ s^-1^, highlighting the faster and stronger quenching response of A116 compared with the wild type.

NPQ was measured with different actinic lights: the mutant confirmed a higher NPQ than the control with all the actinic lights between 150 and 1200 μmol photons m^-2^ s^-1^. At the lowest actinic light herein tested (50 μmol photons m^-2^ s^-1^) NPQ was essentially not induced in either genotype (Supplementary Fig. S2). Across all actinic lights tested, the mutant also showed faster quenching kinetics, with a faster rise during the first minute of illumination (Supplementary Fig. S2). Longer exposure to actinic light (300, 600, and 1200 μmol photons m^-2^ s^-1^) was therefore investigated to determine whether prolonged light stress could induce similar NPQ levels in the two genotypes (Supplementary Fig. S3). Even after 40 minutes of illumination, the A116 mutant maintained higher NPQ levels than the WT. Considering the recovery in the dark of the NPQ kinetics, the NPQ component that increased in the mutant was the fastest-relaxing one, qE. Because qE activation depends on the generation of the proton gradient across the thylakoid membrane, the stronger and faster NPQ induction observed in the A116 mutant could be related to differences in lumen acidification. qE activation can also be monitored by directly acidifying the lumen through the addition of an acid solution to the cells in the medium (Cazzaniga *et al*., 2023; Tian *et al*., 2019); this approach allows overcoming the light dependence of proton gradient generation and evaluating its role in the mechanism of quenching. Upon addition of acetic acid, chlorophyll fluorescence in *H. lacustris* decreased within a few seconds and almost fully recovered after titration of the pH back with triethylamine (Supplementary Fig. S4). A116 showed a similar behavior, but with a stronger decrease of fluorescence. Calculated NPQ indicated that, even under acidification, A116 exhibited higher quenching levels.

### Photosynthetic activity and photosynthetic electron flow of A116 mutant

To evaluate the effect of increased NPQ on the photosynthetic activity, the main photosynthetic parameters as Fv/Fm, PSII operating efficiency (ΦPSII), relative electron transport rate (ETR) and photochemical quenching (1-qL) were investigated in WT and A116 cells, grown in BG-11 at low light (Fig. 2) (Baker, 2008). The light-saturating pulse in dark-adapted cells revealed an identical PSII maximum quantum efficiency (Fv/Fm) in *A116* and WT (0.79 vs. 0.77, respectively). ΦPSII was similar at all the actinic light tested with the exception of 300 μmol photons m^-2^ s^-1^ (the actinic light at which the NPQ difference is highest) where the mutant showed a lower ΦPSII. ETR was similar for the two genotypes only at low actinic lights, while from 300 μmol photons m^-2^ s^-1^ onwards, the A116 mutant showed a lower ETR compared to WT. The photochemical quenching (1-qL) was similar to the WT but higher in the mutant at the two high actinic lights (600-1200 μmol photons m^-2^ s^-1^). To further investigate the photosynthetic electron flow in the *A116* mutant, the light-saturation curve of photosynthesis was monitored as oxygen evolution (Fig. 2). Consistently with ETR measurements, at the lower lights the oxygen evolution rates were similar in the two genotypes, but with illumination higher than 300 μmol photons m^-2^ s^-1^, the mutant showed a decreased oxygen evolution activity. Accordingly, the mutant reached a Pmax (the light-saturated maximum oxygen evolution rate) 40% lower than the WT. Moreover, the light intensity needed to reach half saturation of the oxygen evolution and the slope of linear increase (the phase when oxygen evolution is proportional to the light intensity) were lower in the mutant than in the WT (Supplementary Table S1) indicating a decreased photochemical efficiency in the A116 mutant in high light conditions. The reduced electron transport rate and lower oxygen evolution could be related to greater energy dissipation in the mutant, driven by higher NPQ. Dark respiration was essentially identical in the two genotypes, suggesting that the mutations in the A116 strain did not affect the mitochondrial electron transport chain. Light-dependent proton transport across thylakoidal membranes can be investigated by measuring electrochromic shift (ECS)-induced carotenoid absorbance changes at 520 nm (Bailleul *et al*., 2010a; Lucker and Kramer, 2013). ECS measurements allow indirect determination of the light-dependent proton motive force (*pmf*), which generates the electrochemical proton gradient and induces a shift in carotenoid absorption. ECS measured with different light intensities results in similar *pmf* between the A116 and the WT (Supplementary Fig. S5). ECS was also measured after adding 3-(3,4-dichlorophenyl)-1,1-dimethylurea (DCMU) to the cell to inhibit the linear electron flow from PSII to PSI: the residual *pmf* is related to the cyclic electron flow (CEF) around cytochrome b6f and PSI. The ECS signal attributed to CEF was higher in the mutant, suggesting that decreased linear electron flow in A116 is compensated by increased CEF to maintain similar *pmf* levels.

**Figure 2.**
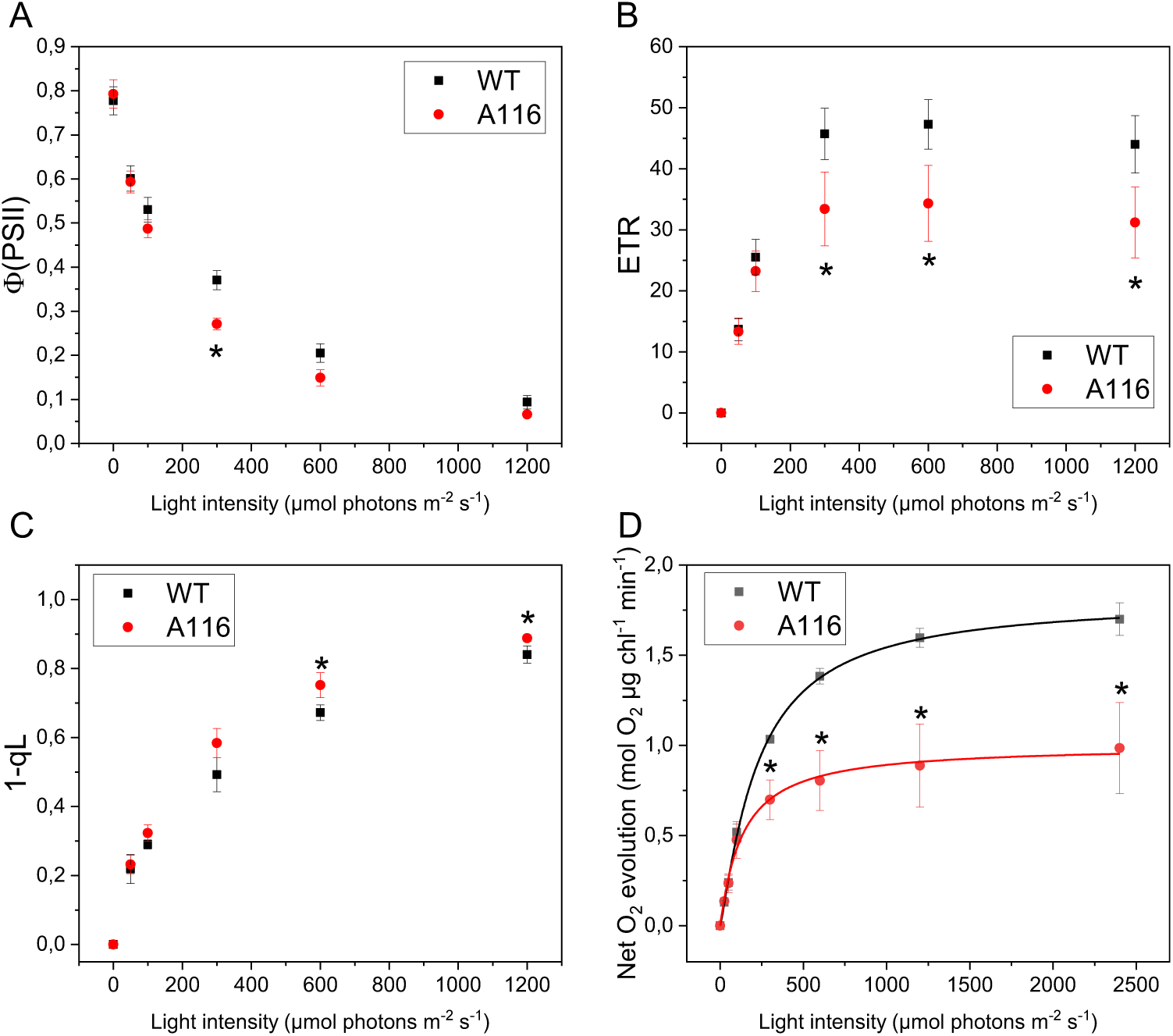
Photosynthetic electron flow and oxygen evolution in A116 mutant. (A) PSII operating efficiency (ΦPSII), (B) relative electron transport rate (ETR), (C) 1 - qL and (D) net photosynthetic O_2_ evolution at different actinic light intensities for wild type (WT, black) and A116 (red). O_2_ evolution data were fitted using a hyperbolic function. Data are expressed as mean ± SD (n > 3). * Indicate significant differences between WT and A116 (P < 0.05). ALT TEXT: Panel A shows PSII operating efficiency in wild type and A116 cells across increasing actinic light intensities, with lower values in A116 at 300 μmol photons m^-2^ s^-1^. Panel B shows relative electron transport rate, which is reduced in A116 at moderate-to-high light. Panel C shows the 1–qL parameter, indicating differences in PSII redox state at high irradiance. Panel D shows net oxygen evolution curves, with A116 reaching a lower light-saturated oxygen evolution rate than the wild type.

### Photosynthetic complexes organization in A116 mutant

Pigments and pigment-binding complexes accumulation were thus investigated in the A116 mutant and compared to the WT, considering that altered chlorophyll content or photosynthetic subunit distribution could significantly influence light-harvesting capacity and NPQ (Cazzaniga *et al*., 2020; Cazzaniga *et al*., 2023; Niyogi *et al*., 1997), As reported in Table 1, WT and A116 strains showed similar chlorophyll content per cell, comparable chlorophyll *a*/*b* ratios, and similar chlorophyll-to-carotenoids ratios. Xanthophylls and β-carotene distributions were further analyzed by high-performance liquid chromatography (HPLC), revealing only minor differences between the two genotypes.

**Table 1.**
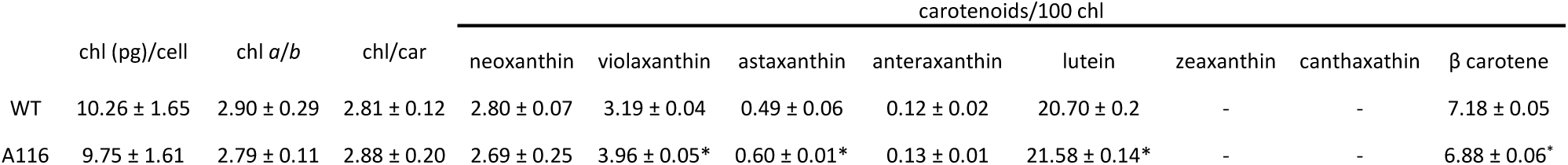
Pigment analysis. Data obtained from cells grown in BG-11 at 25 μmol photons m^−2^ s^−1^. Individual carotenoid contents were normalized to 100 Chls. Data are presented as mean ± SD (*n* = 4). * indicate significant differences between WT and A116 (p < 0.05).

The distribution of photosynthetic subunits was then investigated by separation of the different pigment-binding complexes on native Deriphat-page upon solubilization of thylakoidal membranes α-dodecyl maltoside (Fig. 3A). Based on previous literature, the green bands observed on the gel correspond, from bottom to top, to free pigments (B1), monomeric antenna (B2), trimeric LHCII (B3), PSII core (B4), PSI (B5) and PSI-PSII supercomplexes (B6) (Cazzaniga *et al*., 2014; Mascia *et al*., 2017). The distribution of chlorophyll across the different bands was quantified and was essentially identical in the two genotypes (Fig. 3B). To confirm these assignments, the individual green bands were eluted from the gel and analyzed by low-temperature fluorescence emission spectra (Supplementary Fig. S6). B2 and B3 showed emission peaks at 678-679 nm in accordance with the emission of PSII antenna proteins. B4 showed a peak at 684 nm corresponding to the isolated PSII core. The emission spectra of B5 and B6 were characterized by a peak at 717 nm, indicating the presence of PSI, along with an additional peak at 677-678 nm, likely associated with detached LHC antenna proteins. The organization of photosynthetic complexes was also probed by low-temperature fluorescence emission spectroscopy on whole cells (Fig. 3C). The 77 K spectra showed three main peaks at approximately 685, 696, and 715 nm, corresponding to the PSII antenna, PSII core, and PSI, respectively (Mascia *et al*., 2017). The shape of the spectra were similar in the two genotypes, with maxima at the same wavelengths.

**Figure 3.**
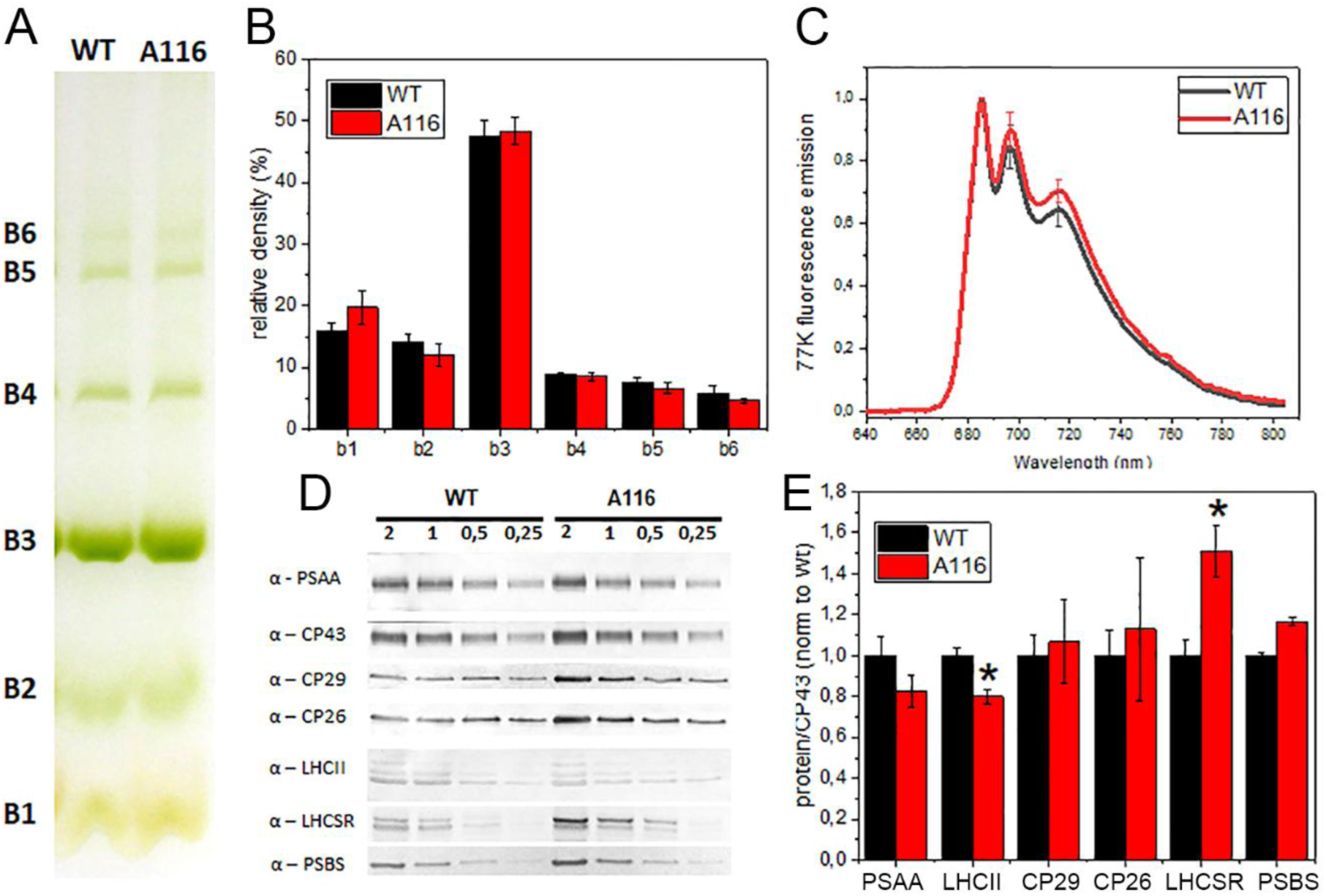
Photosynthetic subunits distribution. (A) Native Deriphat-PAGE of solubilized thylakoid membranes of wild type (WT) and A116. (B) Distribution of chlorophylls among Deriphat-PAGE bands in WT (black) and A116 (gray). (C) Low-temperature (77 K) fluorescence emission spectra of wild type (WT, black) and A116 (red) cells excited at 440 nm. Spectra are normalized to the maximum. (D) Immunotitration of thylakoid proteins in WT (black) and A116 (red) using specific antibodies against PsaA, CP29, CP26, LHCII, LHCSR and PSBS. Lanes correspond to 2, 1, 0.5 and 0.25 μg of chlorophylls loaded per genotype. (E) Densitometric analysis of the immunoblotting results reported in D. Data were normalized to CP43 amount and set to 1 in the case of WT. Data are presented as mean ± SD (n=4). * indicate significant differences between WT and A116 (p < 0.05). ALT TEXT: Panel A shows native Deriphat-PAGE separation of photosynthetic pigment-protein complexes from wild type and A116 thylakoids. Panel B quantifies chlorophyll distribution among the gel bands, showing similar photosynthetic complex organization in both genotypes. Panel C shows 77 K fluorescence emission spectra with comparable PSII and PSI emission peaks in wild type and A116 cells. Panel D shows immunoblots for photosynthetic proteins and NPQ-related proteins. Panel E quantifies protein abundance relative to CP43, showing increased LHCSR accumulation in A116 while most photosynthetic subunits remain similar to the wild type.

The stoichiometry of photosynthetic complexes was also determined by immunotitration using specific antibodies against PSII and PSI subunits (Fig. 3D-E). The PSI/PSII ratio, assessed based on the main core subunits CP43 (PSII) and PsaA (PSI), was similar between the two genotypes. The LHCII/CP43 ratio was slightly decreased (20% decrease with respect to WT). PSII monomeric antenna subunits CP26 and CP29 were present at similar levels in the WT and A116. Overall, these results indicate that the organization of photosynthetic complexes in A116 is comparable to that of the WT and is therefore unlikely to account for the altered NPQ phenotype.

NPQ activation depends on the subunits PSBS and LHCSR, which sense lumen acidification and trigger the quenching mechanism. PSBS is mainly active in plants, whereas LHCSR and LHCSR-like proteins are predominant in microalgae (Li *et al*., 2000a; Peers *et al*., 2009; Troiano *et al*., 2021). Both PSBS and LHCSR genes are also present in the *H. lacustris* genome (Marcolungo *et al*., 2024), and their abundance was quantified by immunotitration using specific antibodies. In A116, PSBS was present at the same level as in the WT, while LHCSR was overexpressed, with the LHCSR/CP43 ratio increased by around 40%. The increased LHCSR content in A116 may therefore contribute to the enhanced NPQ observed in the mutant compared to the WT.

### A116 response under high-light stress

The consequences of increased NPQ induction in A116 compared to the WT were then investigated under high-light conditions. WT and A116 cells were cultivated at low light (25 μmol photons m^-2^ s^-1^) and then shifted at 300 μmol photons m^-2^ s^-1^, the irradiance at which the NPQ phenotype was most evident (Fig. 1). After 48 hours, the responses of the two genotypes was strongly different (Fig. 4). At 25 μmol photons m^-2^ s^-1^, both cultures remained green, and microscopy revealed cells with a uniform green coloration and only minor astaxanthin accumulation, similar in both genotypes. After two days at 300 μmol photons m^-2^ s^-1^, the WT culture drastically shifted toward a reddish color due to strong astaxanthin accumulation, and cells began to aggregate. Bright-field microscopy confirmed that red pigmentation became predominant within the cells. In contrast, A116 cultures maintained a dark green color, and the cells resembled those observed at 25 μmol photons m^-2^ s^-1^. These results indicated that high-light induced transition to haematocysts is impaired in the A116 mutant.

**Figure 4.**
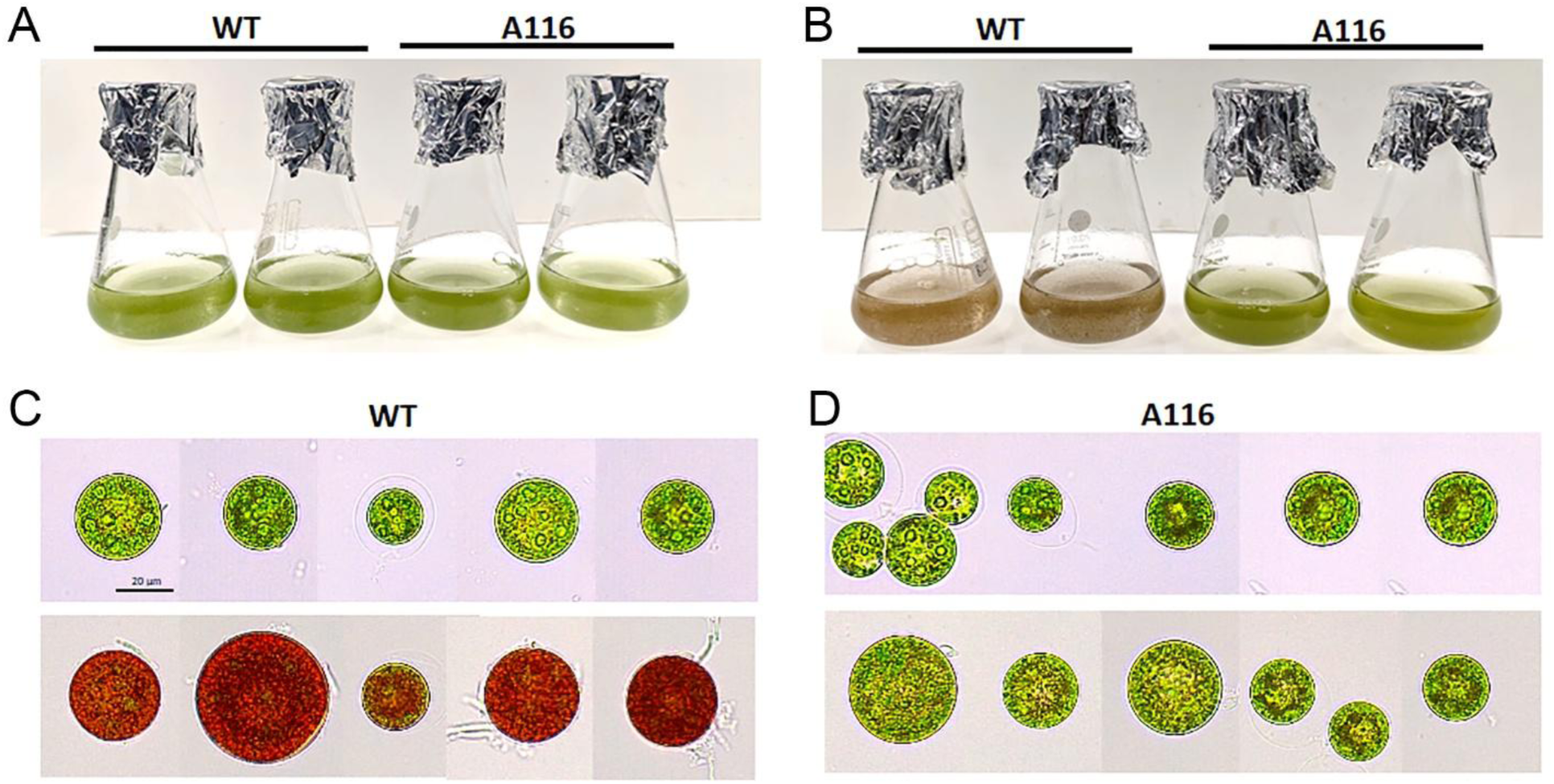
Wild type and A116 shift at higher light. Wild type (WT) and A116 cultures grown in BG-11 adapted at 25 μmol photons m^-2^ s^-1^ (A) or after 48 hours of exposure at 300 μmol photons m^-2^ s^-1^ (B). Bright-field microscopy images of WT (C) and A116 (D) adapted at 25 μmol photons m^-2^ s^-1^ (upper rows) or after 48 hours at 300 μmol photons m^-2^ s^-1^ (lower rows). Scale bar: 20 µm. ALT TEXT: Panel a shows wild type and A116 cultures grown at low light, where both cultures remain green. Panel b shows cultures after 48 hours at high light, where the wild type turns red whereas A116 remains mostly green. Panel c shows bright-field microscopy images of wild type cells before and after high-light exposure, revealing red pigmentation and cell aggregation after stress. Panel d shows A116 cells under the same conditions, with cells largely retaining green vegetative morphology after high-light exposure.

To investigate whether the observed differences were related to NPQ, this photoprotective mechanism was measured in cells shifted to high-light conditions (300 μmol photons m^-2^ s^-1^) for 48 h (Supplementary Fig. S7). After the transition to high light, the maximum NPQ value was reached with faster kinetics compared to the cells grown at lower irradiance. The recovery in the dark was also faster, with a complete NPQ relaxation in two minutes, implying an increase of the qE component with respect to the qI in these samples. The difference between the two genotypes was maintained, and the A116 always showed a higher NPQ at all the lights tested even after the shift. Moreover, in the WT, NPQ induction upon exposure to actinic light was slower than in the mutant and did not reach its maximum in the first minute of illumination. Photosynthetic performance was also assessed in these samples after 24 h of e high-light exposure (Supplementary Fig. S8). In both genotypes, Fv/Fm decreased following the light shift, from 0.77 to 0.54 in the WT and from 0.79 to 0.59 in A116. Similarly, ΦPSII largely mirrored the response observed in low-light–acclimated cells, with comparable values between WT and A116 across most irradiances. The only exception was at 300 μmol photons m^-2^ s^-1^, where the mutant displayed lower quantum efficiency. Consistently, the differences in ETR observed under low-light conditions were no longer evident after high-light acclimation, with A116 showing values similar to WT. The 1−qL parameter was also similar between the two genotypes across the tested light range, although a significantly more reduced PSII state was observed in A116 at 300 μmol photons m^-2^ s^-1^.

### Effects of different abiotic stresses on the transition to the ketocarotenoid synthesis phase in A116

High light is only one of the different abiotic stresses that can induce the transition toward haematocyst and ketocarotenoid producing phase in *H. lacustris*. To assess how ketocarotenoid production responded in WT and A116 cells, the two strains were cultivated under different stress conditions, including nitrogen deprivation, high salinity (1% NaCl), and high-light exposure (Fig. 5).

**Figure 5.**
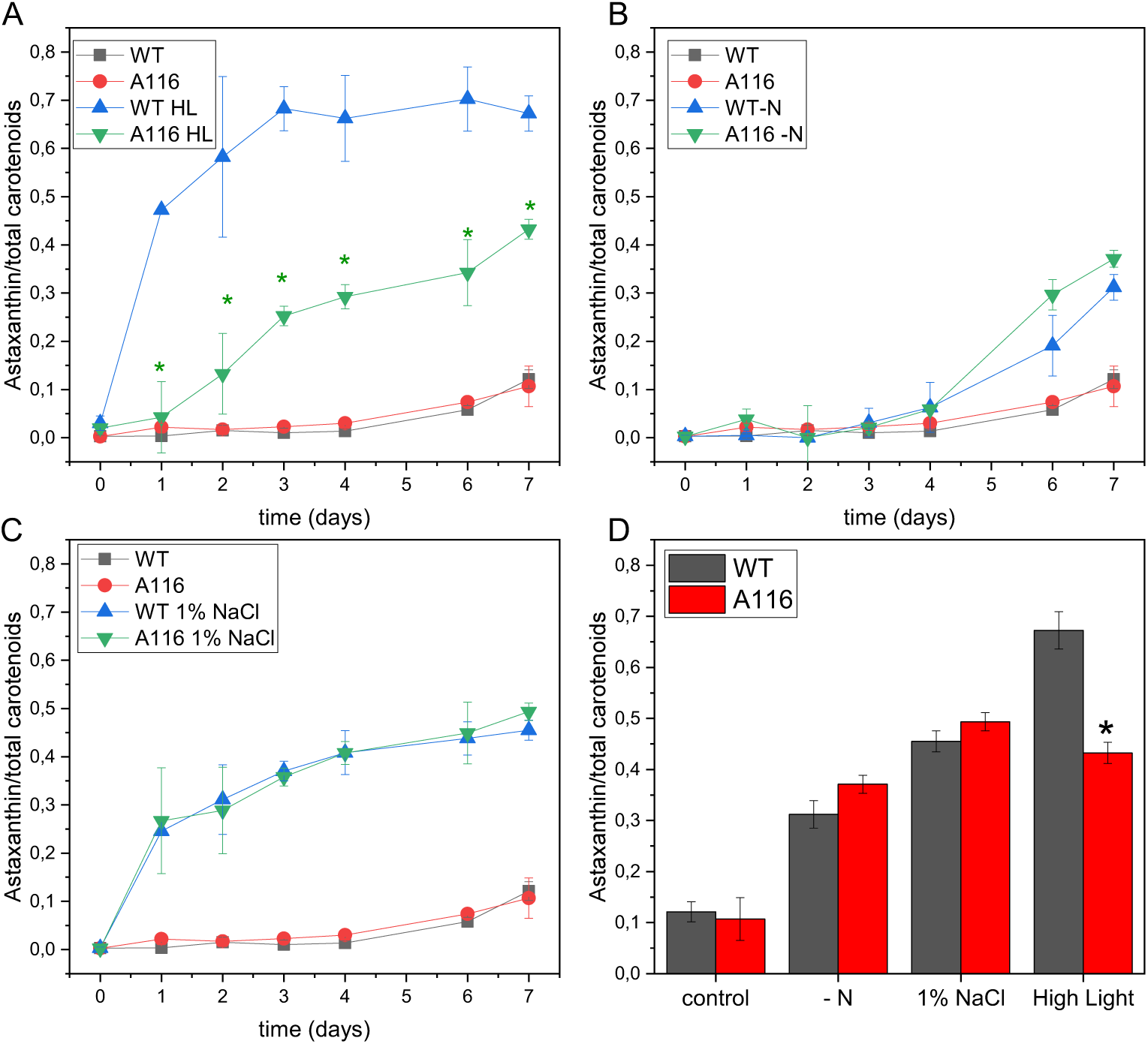
Response of A116 to different stressors. Astaxanthin accumulation in wild type (WT) and A116 Cells were subject to different treatments: (A) high light (300 μmol photons m^-2^ s^-1^, HL), (B) nitrogen deprivation (-N), and (C) high salinity (1% NaCl). (D) Ratio of astaxanthin to total carotenoids measured on day 7. Control samples correspond to WT (black) and A116 (red) cells grown in complete BG-11 medium at 25 μmol photons m^-2^ s^-1^. * indicate significant differences between WT and A116 (p < 0.05). ALT TEXT: Panel A shows astaxanthin accumulation during high-light treatment, with delayed and reduced accumulation in A116 compared with the wild type. Panel B shows astaxanthin accumulation during nitrogen deprivation, where wild type and A116 show similar induction over time. Panel C shows astaxanthin accumulation under high salinity, with comparable responses in both genotypes. Panel D summarizes the ratio of astaxanthin to total carotenoids after 7 days, showing that A116 differs from the wild type mainly under high-light stress.

The transition was monitored as the proportion of astaxanthin relative to total carotenoids. Under high-light conditions, as already observed in Fig. 4, the WT rapidly converted a large fraction of carotenoids into astaxanthin during the first day, reaching approximately 70% of the total carotenoids within three days. A116 showed a delayed response, with ketocarotenoid accumulation occurring only after two days and reached a lower percentage with respect to the control (30% of total carotenoids), confirming the previous results. Nitrogen deprivation showed lower efficiency in inducing astaxanthin accumulation: no ketocarotenoid accumulation was observed during the first three days, and after seven days it reached only ∼30% of total carotenoids. The induction of astaxanthin by nitrogen deprivation in the A116 was similar to that in the WT. Salt stress was more effective than nitrogen deprivation, inducing astaxanthin synthesis from the first day and reaching approximately 50% of total carotenoids by the end of the experiment; accumulation was comparable between WT and A116. These results indicate that astaxanthin production in the A116 mutant is specifically affected by high-light treatment, whereas nitrogen starvation and salinity induce ketocarotenogenesis similarly in A116 and the WT.

### Biomass and astaxanthin productivity

To evaluate the impact of the increased NPQ phenotype observed in A116 on biomass and astaxanthin productivity, WT and A116 cells were grown at high irradiance (300 μmol photons m^-2^ s^-1^) and subjected to repeated dilution cycles (Fig. 6A). Under these conditions, A116 consistently reached higher cell densities than the WT throughout the experiment, resulting in significantly greater biomass accumulation, as confirmed by dry weight measurements (Fig. 6B). A116 cells remained in the vegetative state during light stress and continued to accumulate biomass, while the WT shifted to hematocyst in the first cycle of growth and stopped growing in the following cycles. Consistent with the results reported in Fig. 5, the fraction of astaxanthin per dry weight was lower in A116 than in the WT (Fig. 6C-D). However, due to the higher biomass concentration, the volumetric astaxanthin titer was comparable between the mutant and the WT (Fig. 6E). Because astaxanthin biosynthesis in A116 can be efficiently induced by stress conditions other than high light (Fig. 5), nitrogen starvation was applied at the end of each cycle to trigger the transition to the red stage (Fig. 6F). Under these conditions, both genotypes accumulated comparable levels of astaxanthin on a dry weight basis (Fig. 6G), while the volumetric astaxanthin concentration markedly increased in the mutant, due to the higher biomass accumulation, reaching a final titer of ∼13 mg L^-1^ (Fig. 6H); approximately a 200% increase than in the WT.

**Figure 6.**
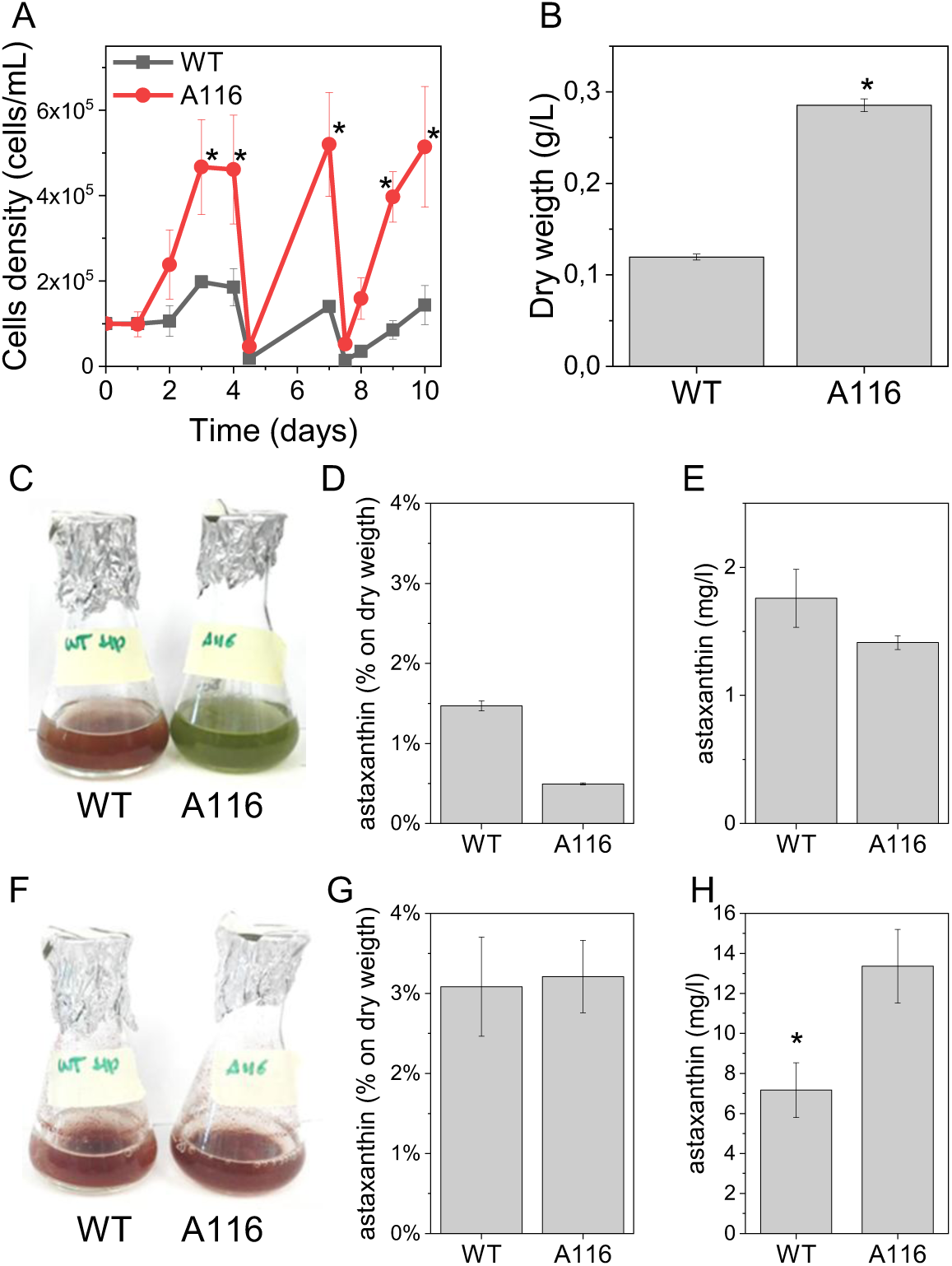
Growth performance and astaxanthin production of WT and A116 during semicontinuous cultivation. WT and A116 cells were grown in flasks under agitation at 300 μmol photons m^-2^ s^-1^ in semicontinuous mode. After 1 day of adaptation, three consecutive growth cycles were performed, with cultures diluted 1:10 every 3 days. (A) Cells density during growth. (B) The average dry weight (g L^-1^) achieved during the growth cycles. (C) Representative images of WT and A116 cells flask at the end of the third growth cycle. (D, E) Average astaxanthin content (% dry weight) and volumetric astaxanthin titer during the growth cycles are shown in (d) and (e), respectively. (F, G, H) Nitrogen starvation was applied at the end of the growth, resuspending cells in BG-11, without N source at 300 μmol photons m^-2^ s^-1^; representative images of WT and A116 cells flask at the end of starvation are shown in (F). The corresponding astaxanthin content (% dry weight) and volumetric astaxanthin titer after nitrogen starvation are shown in (G) and (H), respectively. * indicate significant differences between WT and A116 (p < 0.05). ALT TEXT: Panel A shows cell density during semicontinuous cultivation under high light, with A116 reaching higher densities than the wild type. Panel B shows average dry weight during growth cycles, confirming higher biomass accumulation in A116. Panel C shows representative culture images at the end of the third growth cycle, with wild type cultures redder than A116. Panels D and E show astaxanthin content per dry weight and volumetric astaxanthin titer during growth, respectively. Panel F shows representative cultures after nitrogen starvation. Panels G and H show astaxanthin content and volumetric astaxanthin titer after nitrogen starvation, with A116 reaching higher volumetric astaxanthin production because of its greater biomass accumulation.

In these experiments, cells were grown in flasks without gas exchange and thus in limiting CO_2_; the growth of the A116 mutant was further evaluated under high inorganic carbon availability in controlled conditions using a Multi-Cultivator MC 1000 (PSI) that allows temperature control, OD monitoring at 720 nm and bubbling with air added with 3% of CO2. When cultivated at 300 μmol photons m^-2^ s^-1^ in the presence of 3% CO_2_, the WT exhibited a faster growth rate than A116 (Fig. 7A). Under these conditions, both genotypes showed a reduced NPQ induction relative to cultures grown under carbon-limiting conditions (Supplementary Fig. S9). However, A116 still displayed a faster and stronger NPQ induction than the WT, even at high CO_2_ availability.

**Figure 7.**
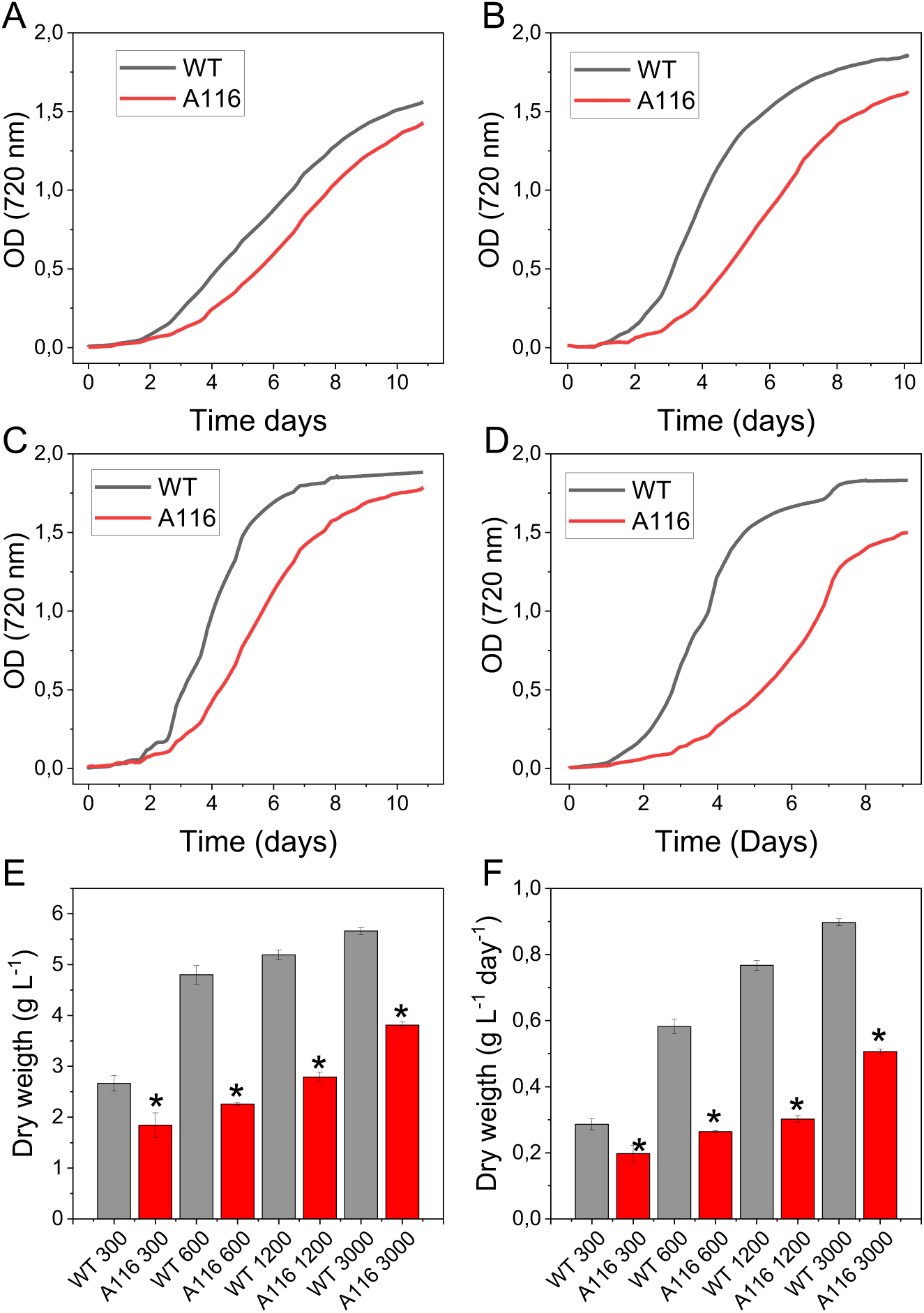
Biomass productivity at different light intensities. Growth curves of wild type (WT black) and A116 (red) lines cultivated with 3% CO_2_ bubbling, 24° C, at 300 (A), 600 (B), 1200 (C), and 3000 (D) μmol photons m^-2^ s^-1^ in BG-11 medium. (E) Dry weight concentration (g L^-1^) of the biomass produced at the end of the growth cycle. (F) Average biomass productivity (g L^-1^ day^-1^) obtained at the growth conditions reported in A-D. ALT TEXT: Panels A to D show growth curves of wild type and A116 cultures grown with 3% CO_2_ under 300, 600, 1200, and 3000 μmol photons m^-2^ s^-1^, respectively. Across all light intensities, the wild type grows faster than A116. Panel E shows final dry weight concentration, which is higher in the wild type at all tested irradiances. Panel F shows biomass productivity, with the wild type reaching the highest productivity at 3000 μmol photons m^-2^ s^-1^ and A116 showing reduced productivity under high carbon availability.

To further assess the effect of light intensity, irradiance was increased to 600, 1200, and 3000 μmol photons m^-2^ s^-1^ under the same conditions (Fig. 7B-D). At all tested irradiances, A116 exhibited slower growth kinetics and lower final biomass accumulation than the WT (Fig. 7E). Biomass productivity increased with irradiance in both strains, reaching a maximum of 0.90 g L^-1^ day^-1^ in the WT at 3000 μmol photons m^-2^ s^-1^. Under the same conditions, A116 achieved significantly lower productivity (0.51 g L^-1^ day^-1^; Fig. 7F). Consistently, astaxanthin production was higher in the WT across all light intensities tested, both in terms of cellular content (% dry weight) and volumetric concentration (Supplementary Fig. S10). In the WT, astaxanthin reached a maximum concentration of 368 mg L^-1^ and a productivity of 58 mg L^-1^ day^-1^, whereas A116 accumulated only 107 mg L^-1^ and 14 mg L^-1^ day^-1^, respectively: approximately one-third of the WT values. Under these conditions, both genotypes showed a reduced NPQ induction compared with the cultures grown under carbon-limiting conditions (Supplementary Fig. S9). However, A116 still displayed faster and stronger NPQ induction than the WT, even at high CO_2_ availability. In these samples, the A116 confirmed an enhanced NPQ mechanism. It showed a fast NPQ rise, reaching the maximum in the first point of illumination and completely relaxing in a few minutes in the dark. At all four light regimes used, the A116 always showed a higher level of quenching than the wild type.

### Identification of the variants in the A116 mutant

The A116 genome was sequenced using Illumina technology and compared to the reference genome (Marcolungo *et al*., 2024) to identify putative variants compared to the background strain. A total of 6973 variants were detected in A116 relative to the WT, including 3179 single nucleotide variants (SNV) and 3794 insertions/deletions (indels). Of these, 71.5% were located within genes, but only 346 mapped to coding regions (CDS), comprising 199 SNVs and 147 indels. Considering the diploid nature of the *H. lacustris* genome (Marcolungo *et al*., 2024), allele distribution was also assessed. Most variants were heterozygous (0/1 genotype), with only three homozygous variants (1/1 genotype) and one heterozygous variant with two alternative alleles (1/2 genotype) relative to the background (Supplementary Table S2 and Supplementary Table S3). Notably, variants affecting both alleles (1/1 or 1/2 genotypes) were not located within the CDS regions and were therefore predicted to have limited effects on protein function (Supplementary Table S3). Among the 346 CDS variants identified in A116, 19 were predicted to have high impact, including stop-gain, start-loss, and frameshift mutations (Supplementary Table S2 and Supplementary Table S3). In total, 17 genes were predicted to be disrupted (Supplementary Table S5), including genes encoding proteins potentially involved in redox metabolism (e.g., FAD-dependent oxidoreductases, flavin reductase-like proteins), protein turnover and signaling (e.g., F-box domain-containing proteins, leucine-rich repeat proteins), and cellular regulation (e.g., methyltransferases, cAMP-related proteins). Additional variants were identified in genes associated with cytoskeletal and flagellar functions (e.g., TPX2, radial spoke protein 3), as well as in several proteins of unknown function. Notably, none of the affected genes have previously been reported to be upregulated under high-light conditions, where the A116 phenotype is most pronounced, with the exception of gene *g1646*, encoding a protein of unknown function (DUF778), which may represent a potential candidate contributing to the observed phenotype, although this interpretation remains highly speculative. The *g1646* gene is predicted to encode a protein with an Armadillo-like fold, a structural motif typically associated with protein–protein interactions and, in several systems, implicated in the regulation of transcriptional and signaling processes (Coates, 2003; Coates *et al*., 2006; Moody *et al*., 2016). It may therefore represent a plausible candidate involved in the regulatory mechanisms underlying the A116 phenotype. All identified mutations were present in a heterozygous state, and no obvious candidate directly linked to NPQ regulation emerged, suggesting that the phenotype may result from indirect effects or the combined contribution of multiple mutations. Additional CDS variants were predicted as missense (186 variants), splice-region (19 variants), or synonymous (129 variants). An exploratory enrichment analysis based on COG functional categories revealed a broad distribution of variant-associated genes across core cellular processes, including replication and repair, energy production, and carbohydrate and lipid metabolism. However, enrichment signals were generally weak, with modest gene ratios and limited statistical support after multiple testing correction, indicating the absence of strongly overrepresented functional categories. Consistent with this, enrichment analyses based on other functional annotation systems (e.g. GO and KEGG) did not yield statistically significant or biologically informative results, likely due to the limited number of genes included in the analysis.

## DISCUSSION

### Increased NPQ phenotype of the A116 mutant

The A116 mutant displays a consistently enhanced NPQ capacity across all tested light conditions, primarily due to stronger and faster activation of the qE component (Fig. 1, Supplementary Fig. S2, Supplementary Fig. S7). Compared to the WT, NPQ in A116 is not only higher in amplitude but also characterized by faster kinetics, indicating a more efficient short-term photoprotective response to fluctuating light.

Although the NPQ photoprotective mechanism is not yet fully understood, several key factors regulating it have been identified. NPQ depends on carotenoid composition, since mutants missing one or more carotenoids or with an altered distribution showed an altered quenching either in plants or microalgae (Niyogi *et al*., 1997; Niyogi *et al*., 1998). However, carotenoid composition in A116 was identical to that of the WT (Table 1), suggesting it does not account for the enhanced NPQ phenotype. NPQ is also affected by the stoichiometry of photosynthetic subunits, particularly within the PSII antenna, which is considered the hypothetical site of quenching. Mutants lacking antenna subunits show pronounced alterations in NPQ kinetics (Dall’Osto *et al*., 2017). In A116, the distribution of photosynthetic complexes, as assessed by native PAGE and low-temperature fluorescence was comparable to that of the WT (Fig. 3). Immunotitration of thylakoid proteins using specific antibodies further confirmed a distribution similar to the WT, with only a minor reduction in the LHCII/CP43 ratio (∼20% less in the mutant). In green microalga *C. reinhardtii*, NPQ is strongly associated with the presence of monomeric antenna subunits CP26 and CP29 (Cazzaniga *et al*., 2020; Cazzaniga *et al*., 2023). The ratio between monomeric subunits and CP43 was similar between the two genotypes. Thus, neither carotenoid composition nor the organization of photosynthetic complexes appears to be altered in A116, suggesting that these factors are unlikely to explain the increased NPQ. Notably, in most previously characterized mutants, alterations in pigment composition or antenna proteins lead to reduced quenching, whereas A116 exhibits the opposite phenotype.

NPQ is triggered by a feedback mechanism involving thylakoid lumen acidification, which arises from saturation of the photosynthetic electron transport chain (Genty *et al*., 1989). Importantly, the enhanced NPQ observed in A116 does not appear to result from differences in lumen acidification. Both prolonged illumination and artificial acidification experiments indicate that A116 retains higher quenching capacity independently of proton gradient formation. Instead, the enhanced quenching is most likely attributable to increased accumulation of LHCSR, the main pH-sensing NPQ effector in green microalgae (Ballottari *et al*., 2016; Camargo *et al*., 2021; Liguori *et al*., 2016; Liguori *et al*., 2013; Peers *et al*., 2009; Steen *et al*., 2022). The higher LHCSR content per PSII observed in A116 (Fig. 3) suggests an increased number of available quenching sites, consistent with the stronger qE component. Altogether, these results point to LHCSR abundance, and not upstream photochemical or bioenergetic differences, as the primary determinant of the altered NPQ phenotype.

### Origin of the A116 mutant and genetic considerations

The stability of the A116 phenotype across multiple experimental conditions, including different light intensities, prolonged illumination, and artificial lumen acidification, supports the conclusion that A116 carries a constitutive alteration affecting NPQ regulation, specifically enhancing the qE component. The persistence of the phenotype even under conditions bypassing light-driven proton gradient formation further suggests that the mutation impacts the intrinsic quenching capacity rather than upstream bioenergetic processes.

The A116 mutant was obtained through an insertional mutagenesis approach based on *Agrobacterium tumefaciens*-mediated transformation, followed by phenotypic screening for altered NPQ (Fig. 1). This strategy enabled the identification of mutant lines with significantly modified photoprotective capacity within a large population of transformants (Fig. 1). Notably, although the initial selection relied on hygromycin resistance, the A116 line lost the resistance cassette, as confirmed by both growth tests and PCR analysis (Supplementary Fig. S1). The stable maintenance of the enhanced NPQ phenotype despite the loss of the selectable marker indicates that the phenotype is not directly associated with the presence of the hygromycin cassette itself. To characterize the genomic alterations associated with the A116 line, whole-genome sequencing was performed, revealing multiple high-impact variants affecting coding sequences. However, establishing a direct genotype–phenotype relationship in *H. lacustris* is intrinsically challenging due to its diploid genome organization (Marcolungo *et al*., 2024) and the heterozygous nature of all identified mutations. The coexistence of multiple alleles, combined with the absence of genetic segregation or complementation analyses, hampers the identification of causative variants among the numerous candidates.

Notably, none of the disrupted genes has been previously associated with NPQ regulation or reported as responsive to high-light conditions, with the exception of gene *g1646* (Supplementary Table S5), encoding a protein of unknown function predicted to contain an Armadillo-like fold. Given the established role of Armadillo-repeat proteins in mediating protein–protein interactions and regulatory processes, this gene may represent a plausible candidate contributing to the altered NPQ phenotype, although its functional role remains to be elucidated.

Overall, these findings underscore both the potential and the limitations of forward genetic approaches in non-model, diploid microalgae. While pinpointing the exact molecular determinant remains challenging, the A116 mutant provides a valuable system to investigate the physiological role of NPQ and its impact on cellular fitness under different environmental conditions.

### Functional consequences of enhanced NPQ

Despite having a similar maximal PSII efficiency in the dark and comparable photochemical parameters under several conditions, the increased NPQ in A116 significantly affects the photosynthetic properties of *H. lacustris* upon exposure to moderate-to-high light (Fig. 2, Supplementary Fig. S8). The enhanced dissipation of absorbed light energy reduces the fraction of excitation energy available for photochemistry, resulting in lower electron transport rates and reduced photosynthetic capacity (Fig. 2, Supplementary Fig. S8). This effect is particularly evident under non-saturating light conditions for the WT, where A116 prematurely activates energy dissipation, effectively limiting carbon fixation. However, under high-light acclimation, these differences become less pronounced, as light is no longer limiting and increased dissipation can instead contribute to photoprotection. In this context, A116 shows reduced photoinhibition compared to WT, highlighting the dual role of NPQ as both a protective and potentially limiting mechanism.

### NPQ modulates the balance between growth and stress-induced differentiation

When *H. lacustris* cells are cultivated under unfavorable conditions for an extended period of time, motile cells lose their flagella and differentiate into palmella cells characterized by thickened cell walls (Boussiba, 2000). Various stress conditions can induce the transition (Zhekisheva *et al*., 2002a). Among the stressors tested, high light represents the most effective trigger, consistent with its strong impact on cellular redox balance and ROS production (Kobayashi, 2003; Scibilia *et al*., 2015; Wang *et al*., 2014; Wang *et al*., 2009).

The delayed transition to the red stage observed in A116 can be explained by its enhanced NPQ capacity. By dissipating excess excitation energy, the mutant reduces over-reduction of the electron transport chain and limits ROS formation, two key signals associated with stress-induced differentiation (Chekanov *et al*., 2016b; Wang *et al*., 2009). As a result, A116 maintains cells in a proliferative state for longer periods, thereby delaying the onset of cyst formation. This shift in the balance between growth and stress response has important consequences: although A116 accumulates lower astaxanthin levels on a per-cell basis during the growth phase, it achieves higher overall biomass, ultimately leading to increased volumetric productivity under certain conditions.

### Context-dependent advantage of increased NPQ

A key outcome of this study is that the effect of enhanced NPQ is strongly dependent on environmental conditions, particularly carbon availability. Under CO_2_-limiting conditions, such as those occurring in poorly mixed flask cultures, photosynthetic electron transport is constrained by limited carbon fixation capacity. This leads to over-reduction of the electron transport chain and increased ROS formation. In this context, the higher NPQ capacity of A116 provides a clear advantage by dissipating excess energy, reducing photodamage, and supporting biomass accumulation. Additionally, the delayed transition to the cyst stage further increases biomass productivity in the A116 mutant.

Under conditions of high CO_2_ availability and efficient mixing, *H. lacustris* enhances the utilization of absorbed light energy for carbon fixation, thereby increasing biomass productivity (Fig. 7). The positive effect of high carbon availability on the biomass productivity of *H. lacustris* is in line with previous studies showing that CO_2_ enrichment enhances growth and alleviates light-induced stress by increasing electron sink capacity and reducing ROS accumulation (Chekanov *et al*., 2017; Wu *et al*., 2020). In this scenario, the enhanced NPQ of A116 becomes detrimental, as it diverts a significant fraction of absorbed energy away from photochemistry. This results in reduced growth rates and lower astaxanthin productivity in the A116 mutant compared to the WT, reflecting an overly dissipative state even under conditions of high carbon availability.

These results demonstrate that NPQ represents a trade-off between photoprotection and photosynthetic efficiency: it is beneficial under stress conditions, where excess energy must be dissipated, but disadvantageous when light energy can be efficiently used for biomass accumulation. The role of carbon availability in regulating NPQ induction under high light has been extensively investigated in *C. reinhardtii*, where high carbon availability leads to downregulation of LHCSR3 expression and consequently reduced NPQ (Arend *et al*., 2023; Maruyama *et al*., 2014; Redekop *et al*., 2022; Ruiz-Sola *et al*., 2023). The primary function of NPQ is to mitigate the formation of chlorophyll triplet states, which increases when photochemical capacity is saturated under conditions of excessive excitation pressure on the photosynthetic apparatus (Niyogi, 1999). Under high carbon availability, the products of the light reactions (ATP and NADPH) are efficiently consumed by carbon fixation, promoting regeneration of ADP, phosphate and NADP^+^ and thereby relieving excitation pressure on the photosynthesis apparatus (Zuliani *et al*., 2024). Under these conditions, the elevated NPQ capacity of A116 is not advantageous for biomass productivity, as it limits photosynthetic performance.

The context-dependent role of NPQ has important implications for the optimization of *H. lacustris* cultivation. Industrial systems often operate under variable and suboptimal conditions, where fluctuations in light and carbon availability can constrain productivity. In such environments, strains with enhanced NPQ capacity, such as A116, may provide improved robustness and sustained growth. Conversely, in well-controlled photobioreactors with optimized CO_2_ supply and light distribution, minimizing unnecessary energy dissipation may be more advantageous. Under these conditions, strains with lower NPQ or more finely regulated photoprotective responses could achieve higher productivity.

Overall, our results identify NPQ as a key target trait for strain selection and engineering, suggesting that its optimal level should be tuned according to the specific cultivation regime rather than maximized per se.

### Conclusions

This study demonstrates that enhanced NPQ capacity in *H. lacustris* profoundly influences the balance between photoprotection and photosynthetic efficiency, with direct consequences on growth, stress tolerance, and metabolic transitions. The A116 mutant highlights how increased energy dissipation can be advantageous under CO_2_-limiting and stress-prone conditions, yet detrimental when carbon availability enables efficient use of absorbed light. These findings identify NPQ as a key regulatory trait whose optimal level depends on environmental conditions, providing a framework for tailoring strain performance to specific cultivation regimes. Overall, this work underscores the importance of fine-tuning, rather than maximizing, photoprotective mechanisms for improving microalgal productivity.

## SUPPLEMENTARY DATA

Figure S1. PCR analysis of antibiotic resistance cassette insertion.

Figure S2. NPQ at different actinic lights.

Figure S3. NPQ under prolonged illumination.

Figure S4. Acid induced fluorescence quenching.

Figure S5. Electrochromic shift analysis.

Figure S6. Low-temperature fluorescence of isolated bands.

Figure S7. NPQ at different actinic lights in WT and A116 mutant upon 48 hours exposure to high light.

Figure S8. Photosynthetic electron flow after high-light shift.

Figure S9. NPQ at different light intensities in airlift photobioreactors.

Figure S10. Astaxanthin production in conditions of high light and high carbon availability.

Table S1. Respiration rate and oxygen evolution.

Table S2. Summary of variant types, predicted impact classes, and major coding consequences, stratified by genotype.

Table S3. Complete list of variant consequence terms identified in the annotated VCF, stratified by genotype.

Table S4. Comprehensive list of genes carrying variants with eggNOG and original annotations, including inclusion status for functional enrichment analysis.

Table S5. List of genes carrying variants in CDS predicted to have a high impact.

## AUTHOR CONTRIBUTIONS

Conceptualization, M.B.; Methodology, M.B., S.C., M.R., M.D.; Investigation, L.G., F.B., N.O., M.M., M.P., E.C., M.R.; Writing – Original Draft, S.C., E.C. and M.B.; Writing – Review & Editing, M.B., S.C.; Resources, M.B.; Supervision, M.B. and S.C.

## DATA AVAILABILITY

All the data described herein are included in Figures or in the Supplemental data. Raw whole-genome resequencing reads are available in the NCBI Sequence Read Archive under BioProject accession PRJNA1450919. Processed datasets, including variant call files (VCF) and genome-wide functional annotation generated using eggNOG-mapper, have been deposited in Zenodo and are accessible at https://zenodo.org/records/21480187.

## ABBREVIATIONS

CDS: coding sequence
CEF: cyclic electron flow
DCMU: 3-(3,4-dichlorophenyl)-1,1-dimethylurea
DMSO: dimethyl sulfoxide
ECS: electrochromic shift
ETR: electron transport rate
GO: Gene Ontology
HPLC: high-performance liquid chromatography
KEGG: Kyoto Encyclopedia of Genes and Genomes
LHCSR: light-harvesting complex stress-related protein
NPQ: non-photochemical quenching
PCR: polymerase chain reaction
PSI: Photosystem I
PSII: Photosystem II
qE: energy-dependent quenching
qI: photoinhibitory quenching
qL: coefficient of photochemical quenching based on the lake model
ROS: reactive oxygen species
SNV: single nucleotide variant
VCF: variant call format
WT: wild type

